# Distinct hippocampal and cortical contributions in the representation of hierarchies

**DOI:** 10.1101/2022.06.15.496057

**Authors:** Robert Scholz, Arno Villringer, Mauricio J.D. Martins

## Abstract

Humans generate complex hierarchies across a variety of domains, including language and music, and this capacity is often associated with activity in inferior frontal gyrus (IFG). Non-human animals have also been shown to represent simple hierarchies in spatial navigation, and human neuroimaging work has implicated the hippocampus in the encoding of items-in-contexts representations, which constitute 2-level hierarchical dependencies. These fields of research use distinct paradigms, leading to disjoint models and precluding adequate cross-species comparisons. In this study, we developed a paradigm to bring together these two areas of research and show that anterior hippocampus and medial prefrontal cortex encode hierarchical context, mimicking findings from animal spatial navigation. Additionally, we replicated classic neurolinguistic findings of 1) left IFG and posterior temporal cortex in the representation of hierarchies and 2) the association between IFG and processing automaticity. We propose that mammals share an evolutionary ancient system for the generation of simple hierarchies which is complemented in humans by additional capacities.

**Highlights:**

- HPC and mPFC activity is specifically modulated by hierarchical context
- Syntax-related regions in the left hemisphere encode for hierarchy in general
- IFGop activity is maintained in later trials for hierarchies but not sequences
- These findings mimic those from animal spatial navigation and neurolinguistics

## Introduction

The ability to generate and process complex hierarchical structures across a variety of domains is a crucial component of human cognition. Some animal species seem able to represent simple hierarchies in social and spatial navigation (Buzsáki & Moser, 2013; McKenzie et al., 2014; Seyfarth & Cheney, 2014). Beyond these basic capacities, humans can generate structures with multiple levels of embedding and across several domains including language, music, and complex action sequencing (Fitch & Martins, 2014). Recent advances on the domains of language, music, and action neuroscience, as well as from comparative cognition, have provided cues on the neural and computational mechanisms underlying the cognitive representation of hierarchies.

In language, the processing of hierarchical syntax relies on two major hubs: inferior frontal gyrus (IFG) and posterior temporal cortex (spanning middle and superior temporal gyri - pMTG and pSTG - and superior temporal sulcus between the two - pSTS) (Friederici, 2011; Hagoort & Indefrey, 2014; Matchin et al., 2017). IFG in particular, has also been implicated in the processing of hierarchies in music and action (Fadiga et al., 2009; Fitch & Martins, 2014), inviting the speculation that this area might be central to the human capacity to process hierarchies in general. Interestingly, IFG has undergone recent expansion along the hominin lineage, both in volume (Buckner & Krienen, 2013) and in its connectivity with other brain areas (Xu et al., 2020) especially with the left pSTS (Rilling et al., 2008).

The role of these areas has been discussed along two rationales: On the one hand, combinatorial *operations* within IFG may be necessary to generate hierarchies (Friederici, 2011; Zaccarella & Friederici, 2015). On the other hand, IFG might implement domain-general *operations* (e.g., relating to working memory, cognitive control or long-term memory retrieval) acting on domain-specific hierarchical *representations* supported by pSTS and other areas (Matchin et al., 2017; W. G. Matchin, 2017; Rogalsky et al., 2011). In support of the latter view, syntactic comprehension seems to activate pSTS already in children below age 7 while IFG only becomes more active at the age of 10 (Skeide et al., 2014) and in adults after extensive training (in comparison with control non-hierarchical tasks) (Jeon & Friederici, 2013). Overall, *relative* activity within IFG seems to increase with processing *automaticity*. Moreover, recent studies which control for the effects of automaticity also highlight the role of pSTS in the processing of hierarchies in the visual and musical domains (Martins et al., 2019, 2020), thus suggesting that pSTS activity is involved in the *representation* of hierarchical structures more broadly, both for low and high levels of automaticity.

A second strand of research has also implicated the hippocampus in the processing of hierarchies across a variety of domains (Berkers et al., 2018; Garvert et al., 2017; Jafarpour et al., 2019; Kepinska et al., 2018; Kumaran et al., 2012, 2016; McKenzie et al., 2014; Schapiro et al., 2013; Stachenfeld et al., 2017; Theves et al., 2016). The hippocampal system – encompassing the hippocampus along with surrounding areas such as the entorhinal cortex - is located within the medial temporal lobe (MTL) and is associated with spatio-temporal cognition (Doeller et al., 2010; Moser et al., 2008), hierarchical planning and navigation (Brunec & Momennejad, 2019), episodic memory (Collin et al., 2015), the processing of sequences and de-novo formation and consolidation of new associative relations (Gridchyn et al., 2020; Kitamura et al., 2017; Klinzing et al., 2019). Furthermore these structures have also been implicated in item-in-context binding (ICB) (McKenzie et al., 2014; Ranganath, 2010) - wherein an ‘item’ is represented as subordinate to a ‘context’ in a simple 2-level hierarchical relationship - and in the formation of schemas through memory generalization processes (Berens & Bird, 2017). Crucially, the capacity to form items-in-contexts representations – in which a contextual cue determines how another cue should be interpreted - is used as a signature of hierarchical cognition in the field of animal spatial navigation and decision making (McKenzie et al., 2014; Ranganath, 2010). Furthermore, a strong link between the hippocampus and the medial prefrontal cortex (mPFC) has been reported in both human (Constantinescu et al., 2016) and animal literature, in which the mPFC has been identified both as contributor of item-in-context information (McKenzie et al., 2016), and as a locus for consolidated long-term memories (Kitamura et al., 2017).

The molecular, physiological and functional properties of MTL circuitry have been mapped extensively in rodents (Diehl et al., 2017; Donato et al., 2017; Marks et al., 2020; Rowland et al., 2016) and two of its properties seem particularly suited to the implementation of hierarchical representations of the kind ‘items-in-context’. First, a common finding in the hippocampus in both animals and humans is that of a functional gradient of spatial and mnemonic granularity (Collin et al., 2015; Strange et al., 2014) along the longitudinal axis of the MTL. In general, the anterior-ventral hippocampus is more active in larger spatial and mnemonic scales – which is more suited for the representation of the global context – and posterior-dorsal hippocampus is more active in finer-grained scales – which is more suited for the representation of local information. This functional organization is also paralleled in structural (Ray & Brecht, 2016) and transcriptomic organization (Vogel et al., 2020). Moreover, waves travelling along the body of the hippocampus – thereby sequentially traversing the different levels of description - have recently been suggested as substrate for multiscale hierarchical planning in the temporal domain (Stachenfeld et al., 2017). Thus, this gradient could potentially be used for the representation of different hierarchical levels, which would be encoded by populations of cells occupying different positions along this gradient. Second, the hippocampal system harbours both place and grid cells whose functional properties can quickly shift from one context to another (Fyhn et al., 2007; Marozzi et al., 2015), a process referred to as *remapping*. This has been shown to enable the formation of context or state-specific memories. These two properties (organization of scale-sensitive representations along a gradient and context/level-specific remapping) could render the MTL suitable to the encoding of hierarchical structures.

Despite the putative function of the hippocampal system in implementing hierarchical ‘items-in-contexts’ relationships, this system is rarely found active in human studies investigating hierarchies in language, music and action. Two reasons could explain this: 1) The hippocampal activation might be specific to the *acquisition* of new item-in-context relations, while most experiments test participants with extensive training. Crucially, item-in-context associations whose acquisition may initially be enabled/facilitated by the hippocampal system could become externalized to the cortex and compressed as part of a consolidation process (Gridchyn et al., 2020; Klinzing et al., 2019; Lehmann et al., 2009; Niethard & Born, 2020; Schwindel & McNaughton, 2011), paralleling changes seen in cortical involvement with increased automaticity. 2) Previous studies usually contrast hierarchical vs. non-hierarchical structures but do not specifically target the contrast between ‘items’ and ‘contexts’ within the hierarchy. Thus, these different strands of literature might focus on different expertise phases and on different aspects of hierarchical processing.

Here, we address these two issues and present two tasks specifically designed to target the contrast between items and contexts in untrained participants. In both tasks, participants are presented with a sequence of two objects, one after the other. Participants are then asked to determine the correct numeric value associated with the specific pair of objects (Figure 1A). The association between object pair and value is defined by one of two rules: hierarchical (HIER) and iterative (ITER), resulting in two otherwise identical tasks. In ITER, the value of the pair is the sum of the individual items’ numeric values, which does not imply any higher-order structure between them (Figure 1C). This is similar to linguistic conjunction in which “[second] and [green] ball” refers the second item in the list [red, ***green***, blue, green]. In HIER, the numeric value of the pair depends on both the composition and order of the items, whereby the first object determines the context for the interpretation of the second. In other words, there is a 2-level set of cues, in which both (context and item) are essential to interpret the pair, but the reference of the second cue (item) is conditional to the context set by the first. This is similar to linguistic subordination, in which the “[second [green]] ball” refers to the fourth item in the list [red, green, blue, ***green***]. This “item-in-context” task structure is also classically used to test for hierarchical encoding of spatial cues in animal spatial navigation (McKenzie et al., 2014; Ranganath, 2010). Participants learn a rule system with a hierarchical decision structure (Figure 1B), which when applied to the interpretation of a sequence of two objects, yields a nested 2-level if-then logical decision task. With this setup we can target not only the general contrast between hierarchy and iteration (“effect of TASK”), but also the contrast between first and second object (“effect of POSITION”). Thus, we can determine whether and how the hierarchical context affects the processing of either object, and isolate hierarchy-level specific contributions.

**Figure 1.**
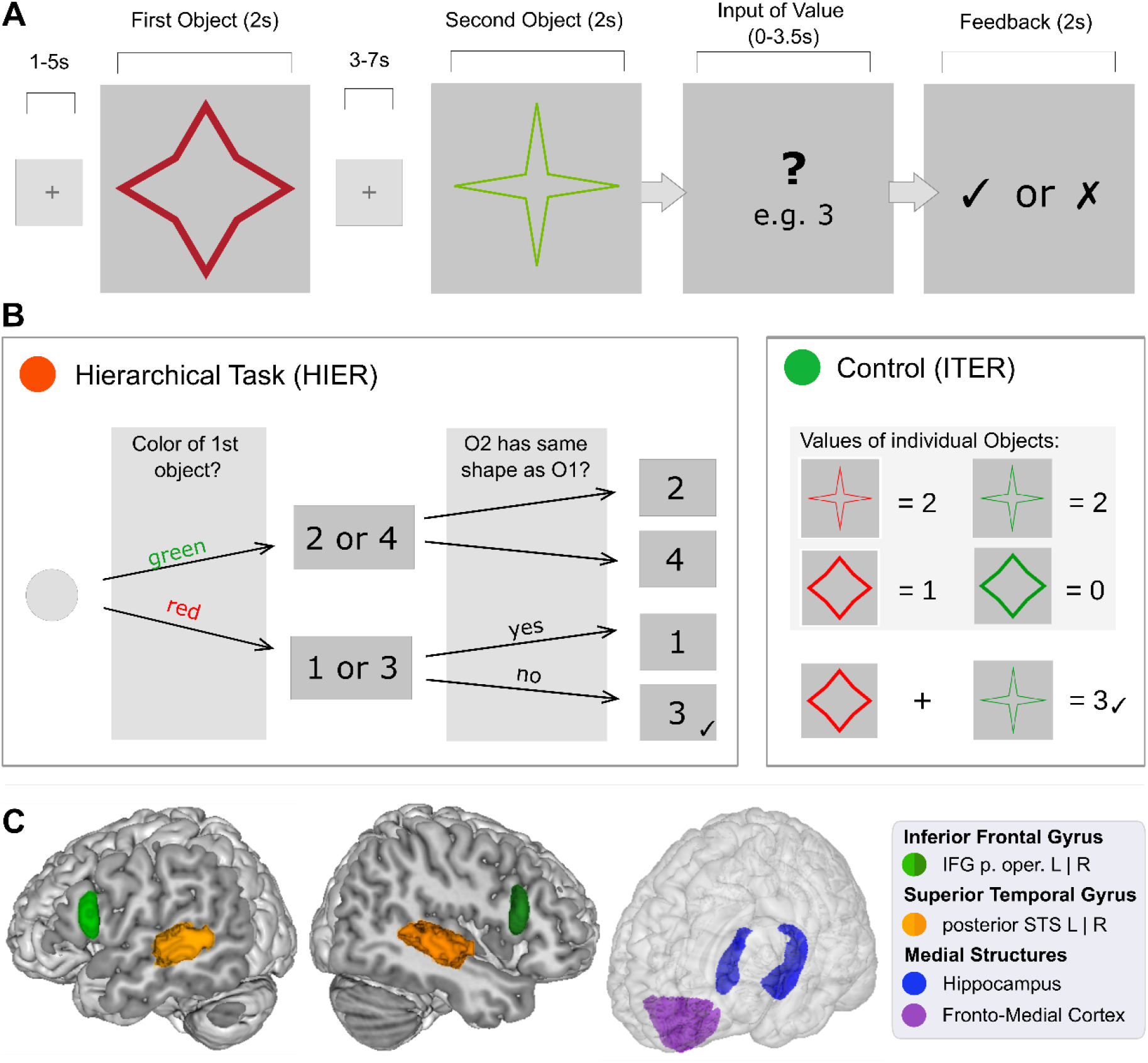
(**A) General Structure of a sample trial of the experimental task.** In this task, participants are presented with two images, sequentially, such that only one image is present on the screen at each time. After the second image is presented, participants are asked to determine the numeric value corresponding to the pair of images and to provide their answer via a button box. Feedback is provided after the answer. Crucially, the numeric value depends on the task condition (see B and C). **(B) Left: Value matrix in the hierarchical task (HIER).** Here, the color of the first object determines the ‘context’ set of possible values ([1, 3] if red and [2, 4] if green). Then the shape category of the second ‘item’ (in relation to the first one) determines the final value of the pair. In this rule system ‘same’ shape does not mean exact visual similarity but rather membership to the same category set (see methods and Figure S3for details). **Right: Value matrix in the iterative task (ITER).** Each object category is associated with a specific value which does not depend on the context. The value of each pair is the sum of the objects’ individual values. **(C) Regions of Interest (ROI) for hypothesis testing ([L]eft and [R]ight).** IFG pars opercularis and fronto-medial cortex masks extracted from Harvard-Oxford probabilistic map (http://neuro.debian.net/pkgs/fsl-harvard-oxford-atlases.html) with 50% and 10% threshold, respectively; Hippocampus mask taken from (Tian et al., 2020) and pSTS mask from (Schaefer et al., 2018). These ROIs are based on regions known to yield activity during hierarchical processing either in human language or animal spatial navigation (see literature review in the text).

Using this design, we aim at clarifying the roles of the hippocampus, IFG, pSTS and mPFC (Figure 1C) in the representation and processing of hierarchical structures – as they have been implicated in hierarchical processing in the research strands reviewed above. Based on this literature, we predict that: 1) The hippocampus will be active in the processing of hierarchical relations (in addition to IFG and pSTS), and more specifically show a differential encoding for the first and the second object (items-in-contexts); 2) different levels of hierarchical organization (first and second object) will be preferentially represented in distinct topographic regions along the hippocampal posterior-to-anterior axis. 3) Given its strong link to the hippocampus, we predict that the mPFC will also be active in the processing of hierarchies, and this activity to increase with automaticity (proxied by experience on the task). 4) IFG and pSTS will be active in the processing of hierarchies in general, but hierarchy-specific activity within IFG – in comparison with a non-hierarchical control task - will increase with processing automaticity. Initially, we also sought to test whether items and context would elicit differential grid-cell activity along the hippocampus axis (like in hypothesis 2). However, our individual stimuli did not elicit robust grid-like activity, precluding testing of this hypothesis (see details of the analysis and results in Supplementary Materials).

## Results

On each trial, two objects were presented, one after the other. After the presentation of Object 2, participants provided the corresponding numeric value of the object pair (Figure 1A). The object pair could be related by a hierarchical (HIER) or an iterative (ITER) rule (Figure 1B and 1C). With this design we were able to assess the specific contributions of TASK ([H]IER vs. [I]TER) and object POSITION ([1]^st^ and [2]^nd^). Crucially, objects and presentation duration were identical across conditions, thus differences in BOLD response could not be explained by simple visual processing. Furthermore, the experiment unfolded across 6 blocks, 3 per rule, in a counterbalanced order (HIHIHI and IHIHIH). By comparing [E]arly and [L]ate blocks (1-3 vs. 4-6) we were able to assess the specific effect of EXPERIENCE with each task. Thus, we focused on the main effects of TASK, and the interactions of TASK x POSITION (H1, H2, I1, I2) [first model] and of TASK x EXPERIENCE (HL, HE, IE, IL) [second model].

### Experimental tasks isolate hierarchical processing and participants’ performance was adequate

Prior to the fMRI experiment, we externally validated the cognitive constructs underlying our tasks (see Supplementary Materials for details). In addition to HIER and ITER, participants performed a Visual Recursion Task (VRT), known to require hierarchical processing, and a Visual Iteration Task (VIT), which does not (Martins et al., 2014, 2019). Previous research has demonstrated the behavioral and neural similarity between VRT and higher-order syntactic processing in language (Martins et al., 2019). In the current study, we found that accuracy in HIER was more correlated with VRT, and ITER more correlated with VIT (see Supplementary Figure S1).

28 new participants completed the fMRI experiment, encompassing 192 trails in total, that is 96 per task. The percentage of correct trials was high for both HIER (*M* = 69.1%, *SD* = 14.4) and ITER (*M* = 68%, *SD* = 14.7) (vs. 25% chance level) and there were no significant differences between the two tasks (*F(27*)=1.22, *p*=.28). We found an interaction between task and block (*F(27)*=1.22, *η* = 0.02, *p*=.02), but no significant pairwise comparisons (*p*<.05, with Tukey correction) between task blocks (means of 66.4, 71.6 and 69.2 for HIER and 69.1, 65.3 and 69.6 for ITER). Finally, we found an effect of block on response time – overall responses became faster in later blocks - (*F(54)*=18.92, *η* = 0.19, *p*<.001), but neither a significant effect of task (*F(27)=0.04, p=.8*) nor a significant interaction of task x block (*F(54)*=0.04, *p*=.95).

### Hierarchical processing activates left prefrontal and posterior temporal cortices

To address the question of which brain structures subserve the processing of hierarchies, we examined the overall effect of TASK (HIER vs. ITER; Figure 2 and Table 1). We found a significant left hemispheric cluster predominantly localized in middle frontal gyrus (MFG), and a second cluster in left temporal cortex, spanning angular gyrus, posterior superior temporal sulcus (pSTS), middle temporal regions and supramarginal gyrus. All reported activations were significant at the cluster level FWE-corrected threshold of *p* <.05. We did not find significant activations for the inverse contrast (ITER> HIER). This shows that hierarchical processing draws on additional cortical resources in the left hemisphere (left MFG and pSTS) when compared to iteration. These results are in agreement with our hypothesis-driven Region of Interest (ROI) analysis in left IFGop (*F(81)=*5.32, *p*=.024) and in left pSTS (*F(81)=*5.27, *p*=.024). For all ROI analyses, we report uncorrected p-values. There was no effect of task in the hippocampus (*F(81)=*1.08, *p*=.303) or mPFC (*F(81)=0.05, p*=.832).

**Figure 2:**
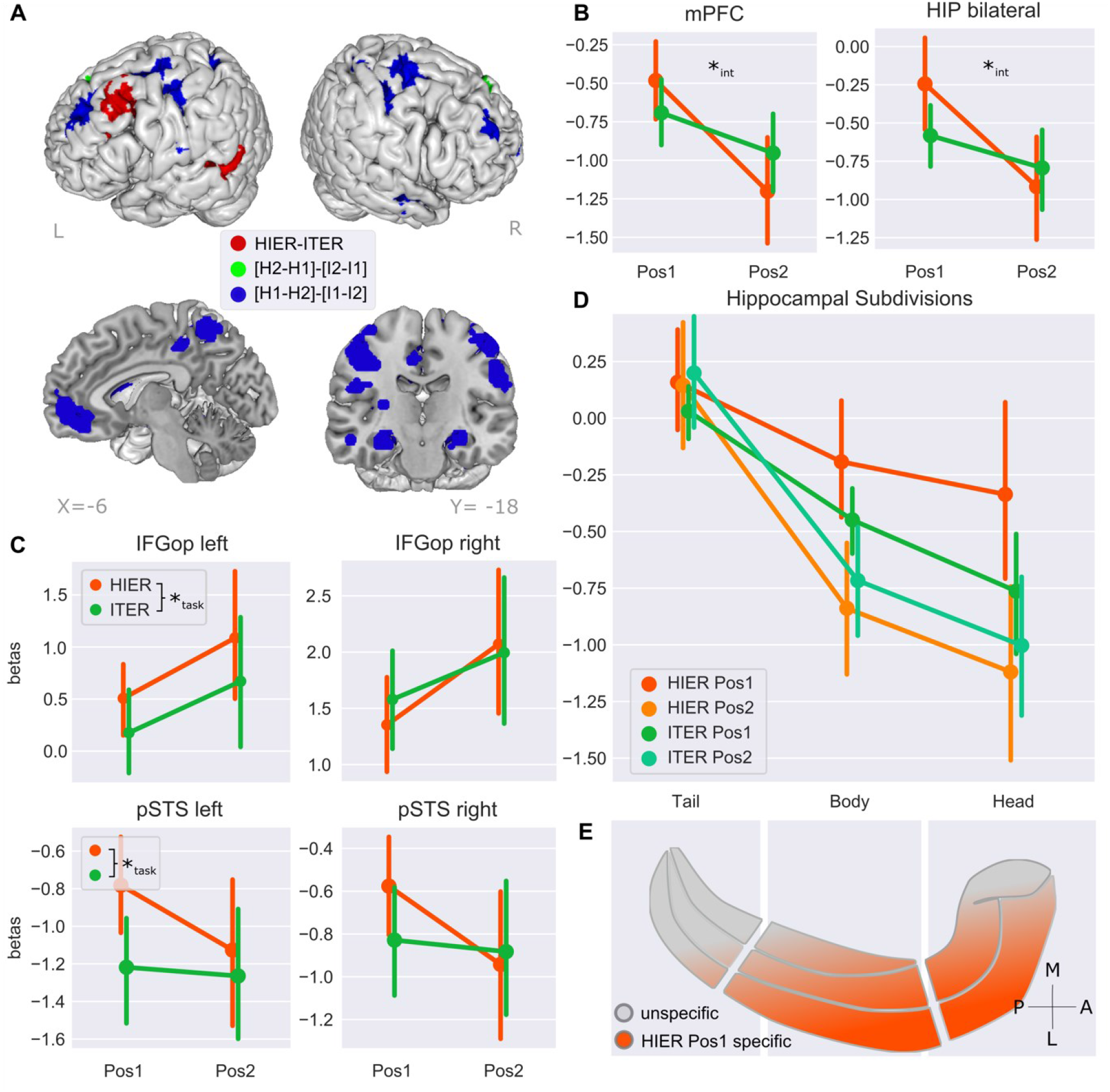
**(A) Whole brain-analysis.** Main effect of TASK (HIER-ITER, red) and interaction TASK x POSITION ([H2-H1]-[I2-I1], green; [H1-H2]-[I1-I2], blue). Statistical t-maps were thresholded and binarized at the cluster level using an FWE-corrected p-threshold of 0.05 and projected on a brain mesh using Mango. **(B, C, D) ROI analyses.** An asterisk indicates a significant effect of TASK (*task) or of the interaction TASK x POSITION (*int; both *p*<.05). (B) Hippocampus (HIP) and medial Prefrontal Cortex (mPFC). (C) Inferior Frontal Gyrus pars opercularis (IFG op) and posterior Superior Temporal Sulcus (pSTS). (D) hippocampus sub-divisions along its long axis. **(E) Schematic POSITION x TASK effect within the hippocampus.** In all subplots, the y-axis describes the average height of beta values for a given condition. For ROI-plots with subject-level datapoints see Supplementary Figure S3.

**Table 1.** Summary of whole brain results for the main effect of TASK and interaction TASK x POSITION. All activations are significant at cluster-level FWE-corrected *p* <.05, and voxel-level at *p*<.001. Cluster and main peaks were labeled using the AAL3 toolbox. In this summary, we only include activations that make up > 5% of a cluster and of the labeled area. **Abbreviations:** cn – cluster number, k – number of voxels in cluster %c – percentage of the cluster belonging to the area, nv – number of voxels of that cluster within the respective area, %r percentage of region occupied by voxels from the cluster.

| Area | H | cn | K | %c | nvcr | %r | x | y | z | Z-score |
| --- | --- | --- | --- | --- | --- | --- | --- | --- | --- | --- |
| **Hierarchy vs. Iteration** |  |  |  |  |  |  |  |  |  |  |
| Frontal Mid 2 | L | 1 | 119 | 85 | 101 | 8 | -45 | 14 | 45 | 4.48 |
| Temporal Mid | L | 2 | 169 | 80 | 136 | 10 | -63 | -40 | 0 | 3.37 |
| Angular | L |  | 169 | 14 | 23 | 7 |  |  |  |  |
| **Interaction 1: Task x Position (Pos2 vs. Pos1)** |  |  |  |  |  |  |  |  |  |  |
| *[H2-H1]-[I2-I1]* |  |  |  |  |  |  |  |  |  |  |
| Frontal Sup Medial | L | 1 | 209 | 60 | 126 | 15 | 0 | 35 | 45 | 5.62 |
|  | R |  | 209 | 24 | 51 | 9 |  |  |  |  |
| **Interaction 2: Task x Position (Pos1 vs. Pos2)** |  |  |  |  |  |  |  |  |  |  |
| *[H1-H2]-[I1-I2]* |  |  |  |  |  |  |  |  |  |  |
| Postcentral | R | 1 | 468 | 29 | 138 | 13 | 33 | -7 | 61 | 5.71 |
| Precentral | R |  | 468 | 27 | 126 | 13 |  |  |  |  |
| Frontal Sup 2 | R |  | 468 | 19 | 90 | 6 |  |  |  |  |
| Frontal Mid 2 | L | 2 | 311 | 60 | 187 | 15 | -30 | 35 | 32 | 5.13 |
| Frontal Sup 2 | L |  | 311 | 37 | 116 | 9 |  |  |  |  |
| Frontal Med Orb | R | 3 | 978 | 12 | 113 | 48 | 30 | 32 | 26 | 5.10 |
| Frontal Mid 2 | R |  | 978 | 11 | 106 | 8 |  |  |  |  |
| Putamen | R |  | 978 | 10 | 96 | 33 |  |  |  |  |
| Frontal Med Orb | L |  | 978 | 8 | 83 | 42 |  |  |  |  |
| Caudate | R |  | 978 | 7 | 72 | 30 |  |  |  |  |
| Putamen | L | 4 | 417 | 32 | 134 | 48 | -21 | 11 | 4 | 5.06 |
| Hippocampus | L |  | 417 | 20 | 82 | 32 |  |  |  |  |
| Caudate | L |  | 417 | 11 | 47 | 21 |  |  |  |  |
| Amygdala | L |  | 417 | 6 | 25 | 41 |  |  |  |  |
| Postcentral | R | 5 | 74 | 74 | 55 | 5 | 27 | -43 | 64 | 4.76 |
| Cingulate Mid | R | 6 | 82 | 57 | 47 | 8 | 9 | -28 | 36 | 4.69 |
| Cingulate Mid | L |  | 82 | 33 | 27 | 5 |  |  |  |  |
| Hippocampus | R | 7 | 84 | 69 | 58 | 22 | 30 | -16 | -19 | 4.68 |
| Precuneus | L | 8 | 206 | 50 | 104 | 11 | -24 | -49 | 64 | 4.58 |
| Parietal Sup | L |  | 206 | 31 | 64 | 11 |  |  |  |  |
| Postcentral | L |  | 206 | 5 | 11 | 1 |  |  |  |  |
| Cerebelum 6 | L | 9 | 70 | 71 | 50 | 11 | -27 | -49 | -28 | 4.53 |
| Precentral | L | 10 | 309 | 38 | 117 | 12 | -24 | -13 | 58 | 4.45 |
| Postcentral | L |  | 309 | 26 | 79 | 7 |  |  |  |  |
| SupraMarginal | L |  | 309 | 6 | 18 | 5 |  |  |  |  |
| Temporal Mid | R | 11 | 116 | 65 | 75 | 6 | 54 | -4 | -25 | 4.31 |

These results cohere with the previous literature highlighting the role of left lateral prefrontal areas and left pSTS in hierarchical processing.

### Hippocampus and mPFC encode hierarchical context

To examine the effect of different object *positions* during the processing of hierarchies (*position 1: ‘*hierarchical context’ vs. *position 2:* ‘embedded item’) we computed the interaction *position* x *task* (Figure 2). When assessing hierarchy-specific contributions during ‘hierarchical context’ vs ‘embedded item’ (i.e. [H1-H2]-[I1-I2]), we find a range of cortical structures and subcortical regions with clusters spanning precentral, postcentral and superior-frontal cortices, medial prefrontal and bilateral orbitofrontal cortices, putamen, caudate, hippocampus, amygdala, bilateral cingulate gyrus, left precuneus, superior parietal and left supramarginal cortex, and right middle temporal cortex (Table 1, interaction 2). Importantly, this set of areas comprises both regions within the dorsal fronto-parietal network, which are related to attention and cognitive effort, but also mPFC, precuneus and hippocampus, which are part of the default-mode network and unrelated to effort *per se*.

Based on our literature review and hypotheses, we performed ROI analyses and confirmed the interaction effect in both bilateral hippocampus (*F(81)*=4.6, *p*=.035) and mPFC (*F(81)*=6.26, *p*=.014). In the hippocampus, this effect is driven by higher betas for the presentation of the first object (H1 > I1) and lower betas for the second object (H2 < I2) when comparing across tasks. When looking at the tasks separately, we find a significant effect of position for HIER (*F(27)*=18.1, *p*<.001), but not for ITER (*F(27)*=2.94, *p*=.098). ROIs with left IFGop (*F(81)*=0.07, *p*=.79), right IFGop (F*(81)*=1.196, p=.277), left pSTS (F*(81)*=1.43, p=.235) and right pSTS (F*(81)*=2.208, p=.141) did not show significant interaction. The reverse contrast of ‘embedded item’ vs. ‘hierarchical context’ ([H2-H1]-[I2-I1]), activated a cluster spanning parts of bilateral superior medial frontal cortices (Table 1, interaction 1).

These findings are in line with our hypothesis that the network comprising Hippocampus and mPFC is involved in ‘items-in-contexts’ representations also in sequences of 2-items connected by a logical hierarchical structure.

### Hierarchical context specifically modulates activity in anterior hippocampus

We predicted that the involvement of the hippocampus could differ along its longitudinal axis. To test this hypothesis, we used functional parcellations that subdivided the hippocampus into three separate regions – head, body and tail - spanning this axis (Tian et al., 2020). We found that mean beta values decrease gradually from posterior to anterior hippocampal regions across conditions (Figure 2D). Furthermore, we found a significant interaction of TASK x POSITION in anterior regions – both head (*F(81)*=4.29, *p*=.042) and body (*F(81)*=4.96, *p*=.029) - but not in the tail (*F(81)=*1.36, *p*=0.25). In sum, the effect of hierarchical context representation in the hippocampus is specific to its anterior regions.

### IFG activity is sustained across early and late trials only for the hierarchical task

Finally, in a separate model, we analyzed the effects of EXPERIENCE for HIER when controlling for ITER (contrast [HL-HE] – [IL-IE]) and found increased *relative* involvement of right opercular IFG, left precentral gyrus, right middle temporal gyrus (MTG), superior frontal gyrus (SFG) and supplementary motor area (Figure 2B, Supplementary Table S2) – in comparison with the control non-hierarchical task. The inverse contrast ([HE-HL]-[IE-IL] did not yield any significant activations. Furthermore, our hypotheses-driven ROI analyses revealed a significant interaction in IFGop bilaterally (left: *F(193)*=4.29, *p*=.04; right: *F(137)*=4.273, *p*=.04). Specifically, while activity was sustained between early and later phases for HIER, it dropped for ITER, thus – similarly to the research reviewed above - the activity of IFGop in the processing of hierarchies increased with experience *relative* to the non-hierarchical control. There were no significant effects for pSTS (left: *F(193)*=.384, p=.536, right: F(193)=0.09, p=.765), hippocampus (F(193) = 0.81, *p*=.396) and mPFC (F(193)=.209, p=.648). Finally, we found a main effect of mPFC increasing activity with experience across both tasks (*F(193)*=4.72, *p*=.031).

**Figure 3.**
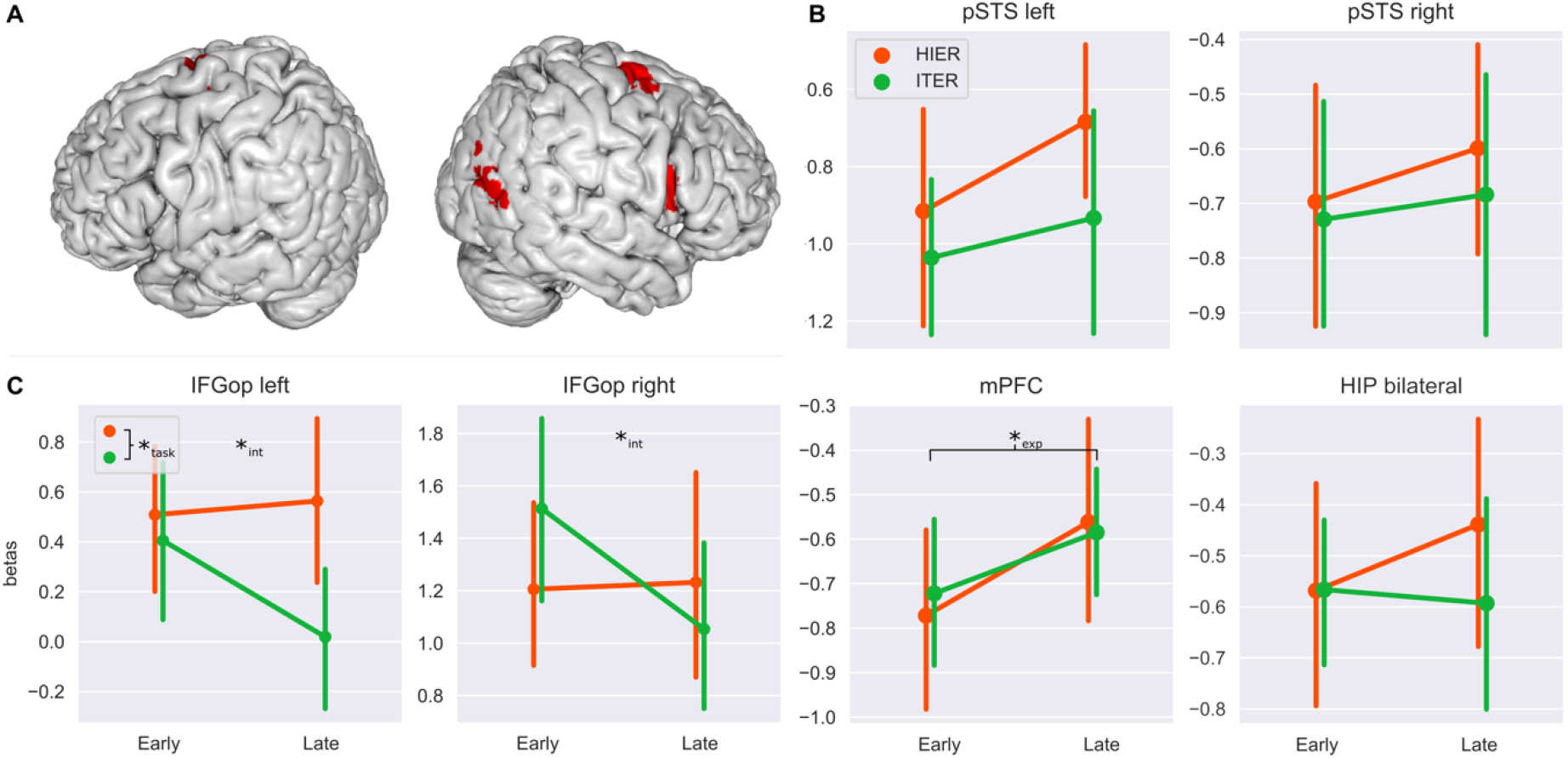
**(A) Whole brain analysis interaction TASK x EXPERIENCE.** Contrast [Hier Late - Hier Early] – [Iter Late - Iter Early] (red). Statistical t-maps were thresholded and binarized at the cluster level using a FWE-corrected p-threshold of 0.05 and projected on a brain mesh using Mango. **(B, C) ROI analyses.** Hippocampus (HIP), right and left Inferior Frontal Gyri (IFG), right and left posterior Superior Temporal Sulcus (pSTS) and medial Prefrontal Cortex (mPFC). An asterisk indicates a significant effect of TASK (*_task_), of EXPERIENCE (*_exp_) or their interaction (*_int_; all *p*<.05).

### Individual stimuli did not elicit grid-like activity, precluding testing of topographic distribution

Using the a similar procedure as previous studies (Collin et al., 2015; Constantinescu et al., 2016), our stimuli varied – and morphed - across two dimensions in the attempt to elicit grid-like activity. Using the GridCat toolbox (Stangl et al., 2017; https://www.nitrc.org/projects/gridcat/), we extracted activity corresponding to 6-fold (grid-like) and 5-fold (control) symmetry and found no differences, meaning that the individual stimuli did not generate robust grid-cell like activity (see details in supplementary materials). Therefore, we did not test for differences across the anterior-posterior axis.

## Discussion

We developed a new experimental paradigm consisting of two tasks which involved hierarchical (HIER) and iterative (ITER) processing. We confirmed behaviorally, that these new tasks correlated specifically with previously validated visuo-spatial recursive and iteration tasks (Martins et al., 2015, 2019), thus showing that they are suitable for segregating hierarchical from non-hierarchy related processing. In fMRI we found that a left lateralized network encompassing lateral prefrontal and posterior temporal cortices was involved in the processing of hierarchies confirming previous studies in the domains of language and vision We also confirmed previous findings demonstrating that IFG activity becomes increasingly specific for the processing of hierarchies (compared to non-hierarchical controls) with increased task automaticity (Jeon et al., 2014; Jeon & Friederici, 2015). Against the backdrop of this behavioral and neuroimaging validation *vis a vis* previous literature and methodologies, our main finding was the involvement of *anterior* hippocampus and mPFC in the processing of hierarchical context, mimicking findings of animal spatial navigation. While mPFC became more active with increased training, hippocampus was equally active in early and late trials.

### Left IFG/MFG and pMTG/pSTS encode hierarchical structure, and hierarchy-related IFG activity becomes more specific with experience

Our ROI results corroborate the involvement of left posterior temporal lobe and IFG in the processing of hierarchies. Whole brain activity peaked in MFG instead of IFG, which could be related to the low degree of automaticity in the processing of our experimental task, compared to the degree of automaticity that is commonly encountered in natural and artificial language experiments, which often draw on already automated processes. Both MFG and IFG seem to be important for hierarchical processing, as lesions in both areas have previously been associated with agrammatic speech (W. Matchin et al., 2020). MFG has been specifically implicated in episodic control in second language acquisition but not first language (Jeon & Friederici, 2013), which supports such a role for the MFG in less automatized contexts. Functional connectivity between left IFG and MFG increases in response to mastery of complex grammar rules (Kepinska et al., 2018), which could be reflective of a stronger involvement of IFG with increased automaticity.

Consistent with this interpretation, our results suggest that the relative importance of IFG in the processing of hierarchies (vs. iteration) increases with the degree of automaticity. While bilateral, this interaction seems more robust in the right hemisphere (unlike in language, but similarly to music and action planning) (Bianco et al., 2015, 2016). These results highlight the idea that while the general principles of hierarchical processing might be analogous across domains, the exact neural circuitry recruited might differ across domains (Blank et al., 2014; Fedorenko et al., 2011, 2012), especially with increased automaticity. In terms of neural and behavioral efficiency, it seems reasonable to surmise that automatic processing operates over domain-specific and not domain-general representations. The use of similar but non-overlapping processing regions is compatible with the formulation of domain-general operations interacting with domain-specific representations (W. G. Matchin, 2017; W. G. Matchin et al., 2017). An alternative explanation may be that at least initially, ITER may draw on a wider range of brain areas - including the IFG - when concurrently exploring multiple task processing strategies (or routes), but that IFG is not specifically active for ITER efficient and automatized processing. This could account for the decrease in IFG beta values in case of the ITER, whereas the betas for HIER remain constant. In other words, IFG-specific activity in the processing of hierarchies could result from the inability to offset processing to other brain regions in contrast to the simpler non-hierarchical task.

### Anterior Hippocampus and mPFC are involved in the representation of hierarchical context

Second, we found that the hippocampus and mPFC – among other regions - were more active in the representation of hierarchical context. Importantly, we found increased activity in areas associated with the fronto-parietal and dorsal-attention networks, which may indicate increased attentional demand, saliency, or task difficulty – even though accuracy, response times and self-reported difficulty were equivalent in both tasks. However, Hippocampus and mPFC are usually considered to be part of (or at least strongly linked with) the default mode network, whose activity canonically decreases with increasing task-directed attention and cognitive demands (Smallwood et al., 2021). Furthermore, we neither found a significant relationship between accuracy and brain activity in our tasks, nor a significant interaction between task and accuracy in explaining neural activity (Supplementary Figure S4). It is thus unlikely that the increased activity of hippocampus and mPFC is explained by attention or cognitive demands alone. Furthermore, we located the effect of hierarchical context to the anterior hippocampus. This is in line with findings showing that global (albeit spatial) context is preferentially represented in ventral/anterior hippocampus. Crucially, such scale-dependent functional organization along the anatomy of the hippocampus might enable the concurrent representation of multiple hierarchy levels in our task, potentially making use of similar organization principles as have been identified previously for both spatial navigation (Buzsáki & Moser, 2013; Kjelstrup et al., 2008; Stensola et al., 2012) and episodic memory (Collin et al., 2015).

### Alternative accounts for the differences across tasks

We believe that the differential task × position effects observed in the hippocampus and mPFC may be related to hierarchical processing, in line with literature reviewed above. For example, upon presentation of the first object, participants need to encode a numeric value in ITER, and a color (in addition to the hierarchical contingency) in HIER. This could potentially explain some of the whole-brain findings, but it seems unlikely that this could also account for the differences seen in the analysed ROIs in the frontal and temporal cortices, as color processing is typically associated with occipital (Kim et al., 2020) and number processing with the parietal cortical regions (Dehaene et al., 1998) instead.

Furthermore, we cannot rule out that there may be some heterogeneity in subjects regarding the employed strategies across tasks (explicit vs implicit processing). Given the high performance and consistently short reaction times across subjects, we would assume that most participants transition to more implicit strategy early on during the experiment. We sought to further ensure this by providing a few training trials for both tasks before the scanning session.

Finally, task difficulty is unlikely to explain our results as we neither find behavioral differences between the tasks nor significant relationships between brain activity patterns and behavioral performance (Supplementary Figure S4).

### Gradual evolution of hierarchical processing on top of pre-existing circuitry

The pattern of activity of mPFC and hippocampus in the representation of hierarchical context and dependencies in our experiment suggests that the underlying mechanisms might be similar to those used in items-in-context representations in non-human animals. Thus, the basic capacity to build hierarchical representations might be available beyond our species. This picture is supported not only by data on spatial navigation and problem solving (McKenzie et al., 2014), but also in the social domain, where for instance, baboons are able to represent several levels of social dominancy based on matrilineal groupings (Seyfarth & Cheney, 2014). Crucially, there is evidence for the involvement of the hippocampus and mPFC in the processing of social dominancy both in humans (Kumaran et al., 2012, 2016; Qu et al., 2017) and other mammals (Dwortz et al., 2022; Wang et al., 2011), hinting on the use of the same system in the processing of more complex social hierarchies.

Recent data on the domain of artificial grammar learning also suggest that monkeys are able to acquire simple hierarchical structures although they require more intensive training than human children (Ferrigno et al., 2020). These data suggest that while the basic capacity for hierarchy representation might be present, other factors could limit the scope and depth of those representations in non-human animals. Several such limitations have been proposed, such as a lower capacity for automatization of knowledge (Schreiweis et al., 2014) and for building abstract and symbolic categories (Sablé-Meyer et al., 2021) due to limited neural supply and connectivity (Changeux et al., 2021). It is possible that IFG and pSTS are more specifically involved in the automatic retrieval of abstract representations than in hierarchical generativity per se, even though the two functions might feedback on each other. More generally, the human capacity to generate complex hierarchies might result from the incremental evolution of brain structures and associated cognitive abilities on top of preexisting circuits (Karmiloff-Smith, 2015) or a recycling of such pre-existing circuitry (Dehaene & Cohen, 2007, 2011).

## Conclusion

We designed a paradigm to bridge findings from separate strands of research, one mainly focusing on language, music and action planning in humans and the other on spatial navigation, item-in-context coding and decision making (especially in animals). Using this paradigm, we have shown that anterior hippocampus and mPFC are more active in the encoding of higher hierarchical levels (hierarchical context), mimicking findings from animal spatial navigation. Additionally, we replicated the classic findings of left IFG and pSTS in the representation of hierarchies, and the increased specific relative role of IFG with higher processing automaticity. With these results we were able to unify research from different domains and species and propose a model for the division of labor. We hypothesize that while mammals might share a system to generate simple hierarchies, this is complemented in humans by additional capacities afforded by an expanded neural circuitry subserving, for example, the automatic processing of abstract representations.

## Materials and Methods

### Participants

We tested 31 healthy right-handed individuals (f=19, mean age=29) recruited from the internal database of the Max Planck Institute for Human Cognitive and Brain Sciences. Of these, the first three participants were excluded from the analysis due to a technical problem which was subsequently fixed. Participants were 18-50 years of age, had normal or corrected to normal color vision, had no history of neurological conditions or extensive use of pharmaceutics/drugs and did not belong to a group of especially vulnerable people (i.e. pregnant or breastfeeding). All participants gave written consent were financially compensated in line with institute regulations. Ethical approval was granted prior to starting the experiment by the Ethics-Commission of the Medicine Faculty of the Leipzig University with the reference 216/19-ek.

### Imaging / acquisition Data

The experiment was carried out in a 3.0-Tesla Siemens SKYRA magnetic resonance scanner (Siemens AG, Erlangen, Germany) using a 32-radiofrequency-channel head coil. During the four sessions, functional magnetic resonance images were acquired using a T2*-weighted 2D echo planar imaging (EPI) sequence with TE = 24 ms and TR = 500 ms. For each session, we acquired altogether a maximum number of 8000 volumes. Due to a hardcoded limit in maximal volume count at around 4000, the sequence had to be restarted for each participant at least once. This restart happened manually at the middle of the experiment after completion of the 3^rd^ block. In two cases, due to interruptions triggered by the participants, a third restart had to be performed. Volumes were acquired with a square FOV of 204 mm, with 36 interleaved slices of 3.20 mm thickness and 10% gap (3.0×3.0×3.2 mm voxel size) aligned to the AC-PC plane, and a flip angle of 45°. T1-weighted images for anatomical co-registration were selected from the database of the institute.

### Experimental Tasks & Stimuli

The main task (Scholz, 2020) consisted in finding the association between pairs of star-shaped objects and a specific integer value. For example, a red object with wide angles followed by a green object with narrow angles could be associated with the value ‘one’ (Figure 1A). The associated integer value depended on the task rule (HIER vs ITER), and on the category of each object. Object category was first defined by a combination of two features: angle width and line thickness. Taking only a single feature by itself in consideration was by design not sufficient to identify the category accurately. This was done to enable consequent grid analysis on a per object level. The categorical space is depicted in Supplementary Figure S2. In this two-dimensional categorical space, objects in the upper right belonged into one category, and objects in the lower left quadrant belonged to another. In this experiment, we used stimuli that extensively covered the space, apart from the space near the border region, in order to reduce the difficulty of object classification. In addition to being placed in one out of the two halves of the two-dimensional space, objects could be either colored in red or green, yielding a total of four distinct object categories.

In each trial, participants saw two objects presented consecutively on the screen. During the 2s long presentation intervals, each object was morphing into its final shape, which was reached at the offset of the interval and served as the basis for object classification. This again was intended to allow for later grid activity analysis. Participants were asked to provide the corresponding integer value from a range of 1 to 4 for the presented object pair. After providing their answer they received visual feedback. To maximize separability of the BOLD response for different task stages, a fixation period of varying duration was introduced at the beginning of each trial (1-5s) and between both object presentations (3-7s). A sample trial is depicted in Figure 1A. In each trial, the association between objects and value depended on the specific task rule. In HIER, there was a hierarchical dependency between the first and second object (Figure 1B), while for ITER the value of the pair was simply the sum of the values of its objects (Figure 1C). For each participant, the same base set of 96 object pairs was twice shuffled, and each time split into three blocks yielding 3 blocks for each task (HIER and ITER). Those blocks were presented in interspersed order, and the starting task was counterbalanced across participants.

### Procedure

During an out-of-scanner training phase, participants initially learned to categorize the available objects into one of four categories in a forced-choice paradigm. Each trial included a target object that was classified by selecting one of two objects chosen as prototypes of their respective category (one correct and another incorrect). Overall, participants had to successfully complete 84 object categorization trials. If the participant did not identify the class of the target object correctly, the trial was pushed to the end of the queue. The training ended once all trials were successfully completed.

Afterwards, participants were explicitly instructed with the rules of both HIER and ITER using a pre-recorded instructional video and a textual summary of the rules. Participants were then asked to solve 8 trials of each before entering the scanner. Additional oral instructions on up to 4 trials per rule were provided by the experimenter if necessary. In the scanner, participants were presented with six alternating blocks - three of each rule. Each block was composed of 32 trials and was preceded by a screen which indicated the task rule. The order of blocks (starting with either HIER or ITER) was counterbalanced across participants. This blocked design was chosen to minimize habituation effects. Participants received visual input through of a dual mirror projection system and could indicate their choices by means of a 4-key button box placed in comfortable reach of their right hand. After scanning, participants completed a short questionnaire. The entire procedure lasted approximately 1.5-2h.

### Quantification and Statistical Analysis

fMRI data were preprocessed and analyzed using statistical parametric mapping (SPM12; Welcome Trust Centre for Neuroimaging; http://www.fil.ion.ucl.ac.uk/spm/software/spm12/). The preprocessing followed standard procedure using the available SPM preprocessing template. This included slice time correction (using cubic spline interpolation), motion correction, and anatomically (T1)-guided and magnetic-field corrected co-registration and spatial normalization of the functional data to standard stereotactic space/Montreal Neurological Institute (MNI) space. Lastly the data was smoothed using a 3D Gaussian kernel with full-width at half-maximum (FWHM) of 6mm.

For single-subject analyses, evoked hemodynamic responses for the different event types (task, object position, feedback type and key presses) were modeled using a single standard general linear model that additionally included six parametric motion correction regressors. A separate single-subject analysis was conducted for the effect of experience, by classifying the respective event onsets into either first (early) and second (late) phases of the experiment. The outputs of the single subject models were used in the group analysis which was conducted using a flexible full factorial design (one GLM for subject, task and position; and a second for subject, task, experience). Using these second level GLMs, we computed whole brain group T-contrasts. FWE-correction of the resulting statistical maps was performed on cluster level and significant clusters were identified using a FWE-corrected p-threshold of 0.05. For the definition of the linear models, t-contrast calculation and cluster correction, the respective standard functions provided by SPM were used.

To test for the involvement of hypothesized regions, we performed region of interest analyses (ROIs) by contrasting mean activity across conditions. The masks for IFG pars opercularis and fronto-medial cortex masks were extracted from the Harvard-Oxford probabilistic atlas (http://neuro.debian.net/pkgs/fsl-harvard-oxford-atlases.html) with 50% and 10% threshold, respectively. Furthermore, the mask for the hippocampus and its subregions were taken from (Tian et al., 2020) and the “LH_Default_Temp_2” parcel from (Schaefer et al., 2018) was used as pSTS mask. We used the REX toolbox (http://web.mit.edu/swg/software.htm) to extract mean single-subject beta values across different conditions and ROIs. Using these Betas, we computed the main effect of TASK and interaction effects of TASK x EXPERIENCE and TASK x POSITION with mixed models (fixed effect omnibus tests) in Jamovi (*The Jamovi Project*, 2021). Brain overlays were plotted using mango (http://ric.uthscsa.edu/mango/).

### Resource Availability

Individual participant behavioral and neuroimaging data cannot be made publicly available. Group contrasts, analysis scripts and the experiments implementation using psychopy have been deposited in an Open Science Framework repository and can be accessed at https://osf.io/56nxh/. The Harvard-Oxford comes with FSL (https://fsl.fmrib.ox.ac.uk/fsl/fslwiki). The Melbourne Subcortex Atlas (Tian et al., 2020) containing the remaining ROIs is available on Github (https://github.com/yetianmed/subcortex). Further materials and code for the data analysis are available upon request to the lead contact, Robert Scholz.

## Acknowledgments

We thank the group of C.F.Doeller for their comments and discussion, Jöran Lepsien for his helpful suggestions on neuroimaging setup and statistical analysis, Simone Wipper for her assistance during participant recruitment and neuroimaging data acquisition and Ramona Menger and Maria Paerisch for their assistance in administrative questions.

## Author contributions

R.S., A.V. and M.J.D.M jointly conceived of the experiment and the hypotheses. R.S. conducted the study, piloting, data collection and analysis. M.J.D.M assisted in the analysis. R.S, M.J.D.M and A.V. wrote the manuscript.

The authors declare no competing interests.

