## Supplementary Information for "Distinct hippocampal and cortical contributions in the representation of hierarchies"

### Validation of Experimental Tasks

Prior to the fMRI study, we conducted a behavioral study with 27 participants to evaluate whether our novel logical hierarchy task tapped into the specific underpinnings of hierarchical processing (Scholz, 2020). In our procedure, participants completed a minimum of 30 trials and a maximum of 98 trials with the logical HIER task and the control ITER task.

In this validation procedure, participants were not explicitly shown the task categories and rules and had to acquire them by trial and error (with feedback). We recorded several measures of performance. First, we measured the number of correct trials within the minimum mandatory set of 30 stimulus (f30). Second, we measured the number of trials required to reach a first criterion of 5 correct trials within a shifting window of 8 trials (*nfc – n to first criterion*). Finally, we also recorded the number of trials to reach a second criterion (*nsc)*: Every time participants passed the first criterion, they were queried with a set of 8 trials, one of each stimuli category. Passing the second criterion required a minimum of 7 correct trials out of 8.

Participants showed similar scores for the acquisition of the ITER (f30: $M = 16.11$, $SD = 3.87$; nfc: $M = 23.75$, $SD = 13.04$; nsc: $M = 43.39$, $SD = 23.81$) and HIER (f30: $M = 16.63$, $SD = 4.02$; nfc: $M = 24.67$, $SD = 10.77$; nsc: $M = 39.89$, $SD = 21.45$), with no significant differences in these scores across tasks: f30 (Wilcox' $V = 145.50$, $p = .66$), nfc (Wilcox' $V = 113.50$, $p = .30$) and nsc (Wilcox' $V = 103.00$, $p = .46$). This shows that participants generally were able to extract the underlying rule in HIER and ITER even with only limited training and instructions beforehand.

For external validation, we assessed the correlation HIER and ITER, and a set of tasks previously shown to tap into hierarchical cognition: the Visual Recursion Task (VRT) and the Embedded Iteration Task (EIT). VRT and EIT measure the ability to represent recursive hierarchical embedding and simple iteration in the visual domain, respectively (Martins et al, 2016). Across all measures, we found that HIER correlated more strongly with VRT ($r_{f30}= 0.38$, $r_{nfc}= -0.33$, $r_{nsc}= -0.26$) than with EIT ($r_{f30}= 0.23$, $r_{nfc}= -0.07$, $r_{nsc}= -0.22$), whereas ITER correlated more strongly with the EIT ($r_{f30}= 0.45$, $r_{nfc}= -0.21$, $r_{nsc}= -0.35$) than with VRT ($r_{f30}= 0.23$, $r_{nfc}= -0.11$, $r_{nsc}= -0.19$). Further linear models corroborate this double dissociation between HIER-VRT and ITER-EIT (Scholz, 2020).

Taken together these results suggest that 1) HIER and ITER are of similar difficulty, which was also confirmed by oral reports, with some subjects finding the hierarchical task harder and others the iterative, 2) HIER is more specific to hierarchy than ITER, 3) ITER can serve as a control task that is more associated with non-hierarchical, iterative processing.

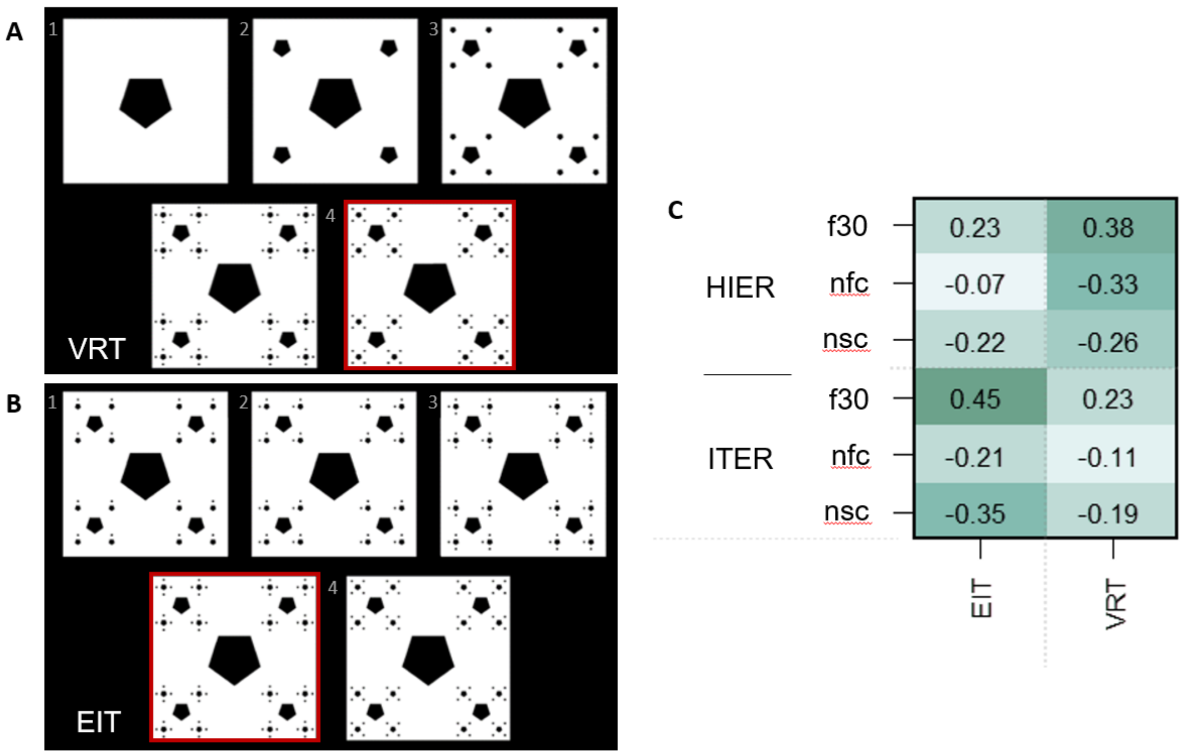


**Figure S1. Validation tasks and cross-task correlations.**

**(A,B)** sample trials of the Visual Recursion Task (VRT) and Embedded Iteration Task (EIT), used to externally validate our novel tasks HIER and ITER. In both VRT and EIT, the upper rows contain the first 3 steps of an iterative process, presented sequentially to the participant. Then, two images are presented simultaneously in the bottom part of the panel, and participants are asked to choose the correct $4^{th}$ step corresponding to the continuation of the first 3 shown above. For an illustration, correct choices are framed in red. **(C)** correlation coefficients (Kendall's W) between our indicators of HIER and ITER performance (f30 – number of correct trials within the first 30; nfc – number of trials until first criterion – 5/8; nsc – number of trials until second criterion – 7/8) and VRT and EIT.

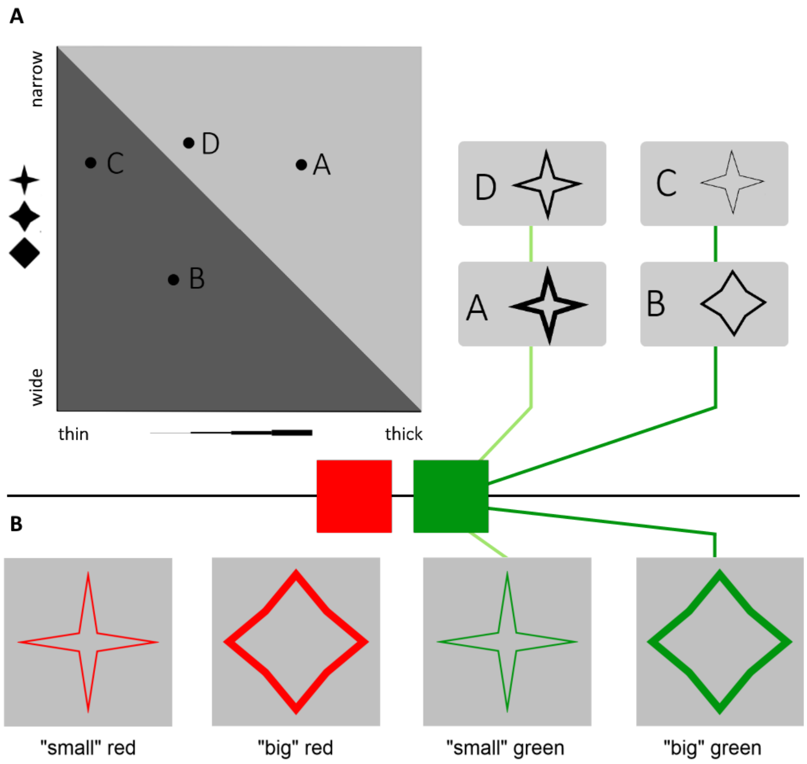


**Figure S2. Assignment of stimuli into four categories.**

**(A)** All objects used in this experiment were mapped along a space of two dimensions: angle width and line thickness. The space was divided into two halves which defined two object categories. For instance, objects in the upper right half (A and D) belonged to one group, and objects in the lower left half (C and B) belonged to another. The objects used as stimuli covered this categorial space extensively. **(B)** In addition to being subdivided according to their angle width and line thickness, objects could be colored in either red or green. Hence, there were four object categories in total.

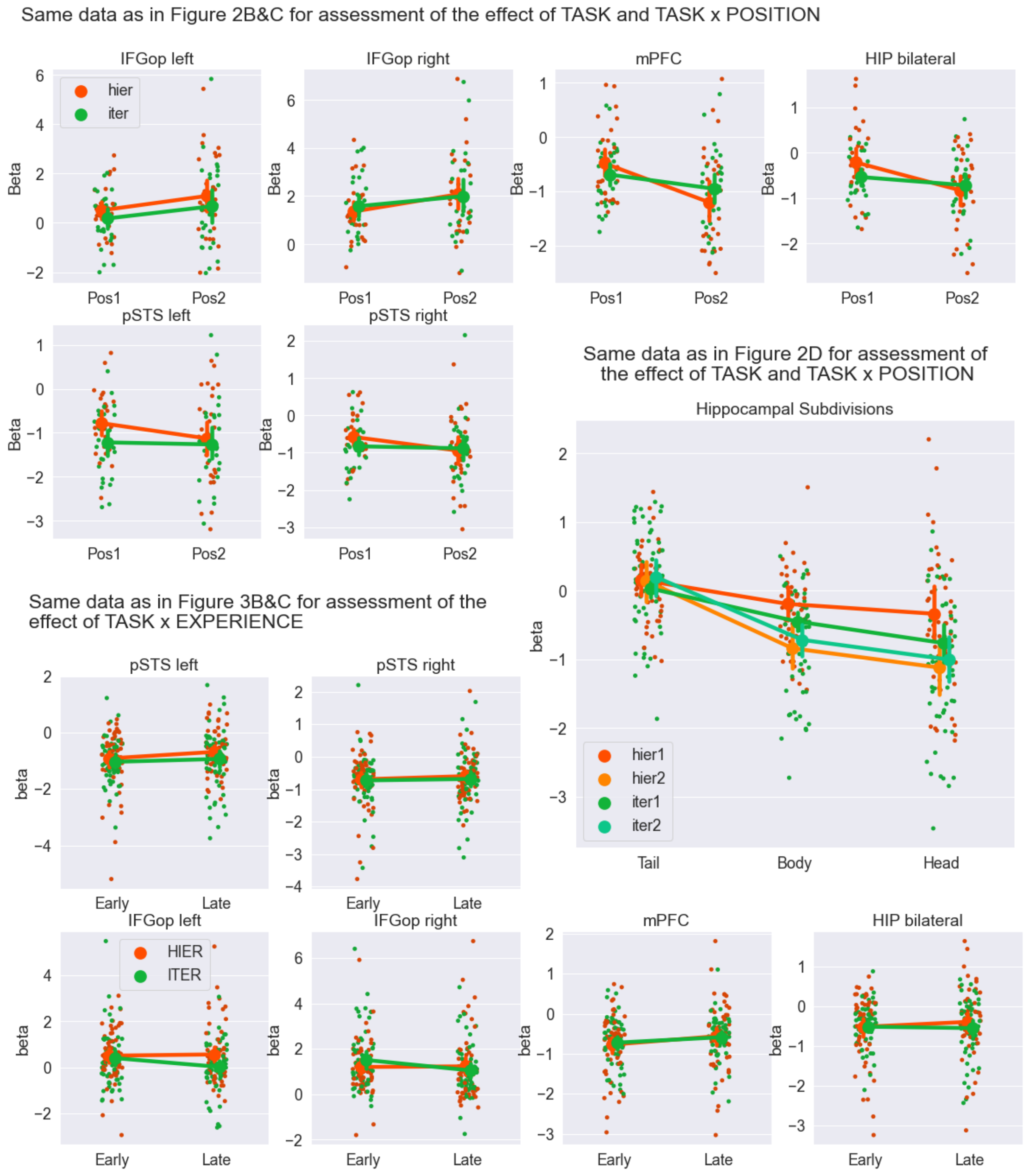


**Figure S3. Scatterplots for all conducted ROI analyses**

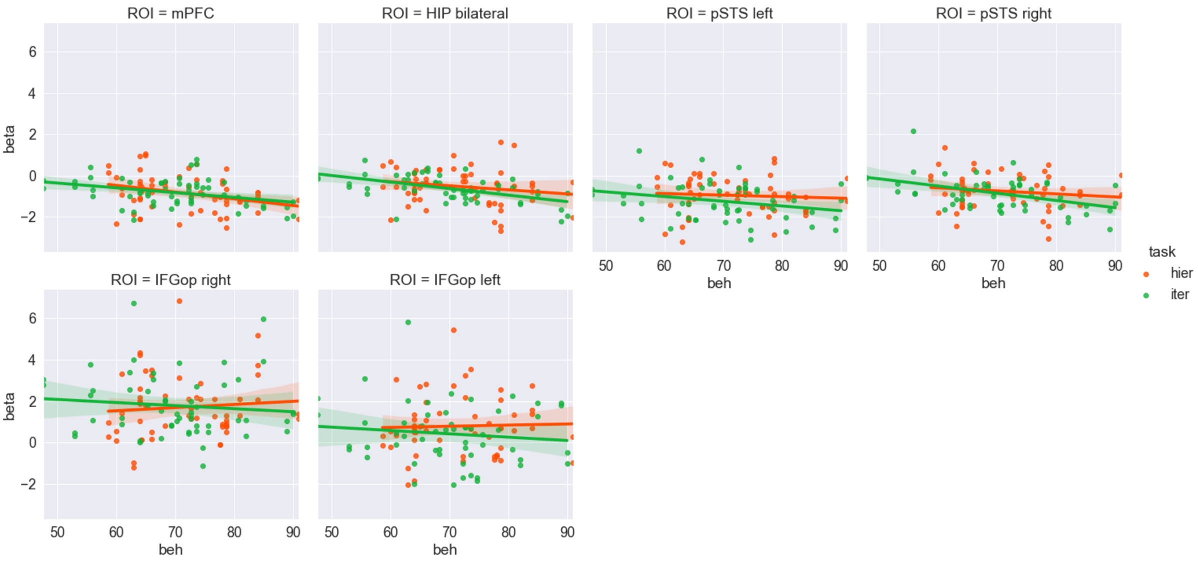


**Figure S4. Relationship between ROI activity (raw betas) and % of correct trials.**

Each data point corresponds to each participant's ROI activity x %correct across all conditions. We ran the following Linear Mixed Models for each ROI: ROI beta = task + %correct + task*%correct + (1|participant). In all cases (ROIs), we found no significant main effects of %correct in ROI activity, and no significant interactions task*%correct (all $p > .05$). In addition to the absence of differences in accuracy and response times in the behavioral data, this activity pattern does not support the idea that differences in difficulty between HIER and ITER are driving HIER > ITER effects.

**Grid Analysis**

In this experiment, we also attempted to stimulate activity of the grid cell system by using conceptual objects which varied and morphed in a 2-dimensional system, as in Collin *et al.* (2015) and Constantinescu *et al.* (2016). Unfortunately, we did not obtain a stable grid pattern of activity. The failure to show grid-cell involvement can have multiple reasons, as stimuli may not have been (perceptually) adequate, the overlap in trial-by-trial BOLD responses may have been too high for parametric analysis, or the analysis could have been compromised by the existence of a third dimension of color, as additional dimensions may have diverse effects on grid cell coding (Marozzi et al., 2015). A failure to demonstrate involvement of grid cells does not necessarily implicate its absence and in the future we will replicate this experiment with more targeted stimuli.

We looked at hexagonal / grid-like modulation of the functional signal during object presentation, which has been found in regions such as the entorhinal cortex and medial PFC (Collin *et al.*, 2015; Constantinescu *et al.*, 2016). In order to allow for such an analysis, the paradigm made use of morphing objects instead of static ones. Two consecutive object morphs of two seconds each, separated by a jittered interval of 3-7 seconds, were presented for each trial. The morphing hereby should mimic a trajectory through a two-dimensional concept space (CS) spanned by the dimensions of line thickness and object shape (ranging from squared to star shaped objects). The participants were asked to base their estimate of object pair values on the morph outcomes (i.e. the final configuration of the object at the end of the trajectory). To derive the correct object classes and therefore be able to estimate the correct object pair value, participants had to closely track each object position in this artificially constructed CS. Given the generally high scores reported in the results section, we can conclude that participants did indeed keep track thereof. The ratio of change between the two parameters (line thickness and form) during the morphing of an object were taken to determine the angle of the trajectory. We predicted that both TASK and POSITION could influence the encoding of the CS and therefore estimated the grid-like modulation individually for different groupings of object presentations (i.e. separately for H1, H2, I1, I2 or HIER and ITER).

We conducted this analysis using the GridCat toolbox (Stangl et al., 2017; https://www.nitrc.org/projects/gridcat/) and also employed a manual approach to crosscheck. In the latter, we estimated the betas for two parametric regressors - $\beta_{sin} \sim sin(XFsym * angle(trial))$ and $\beta_{cos} \sim cos(XFsym * angle(trial))$ - for each voxel in the brain using a GLM in SPM, where *angle(trial)* corresponds to the angle of the trajectory through the concept space throughout each trial. We used an X-fold-symmetry (Xfsym) of 6 for the estimation of a hexagonal grid code and a Xfsym of 5 in the control condition. The grid magnitude GM and grid main orientation GO for each voxel was calculated by $GM = sqrt(\beta_{sin}^2 + \beta_{cos}^2)$ and $GO = arctan2(\beta_{sin}, \beta_{cos}) / Xfsym$ respectively. A similar approach is employed by GridCat. Additionally, each subject's data set was split in half prior to analysis. GM and GO are only estimated on the first half, and then used to predict the activity for aligned vs misaligned trajectories in the second half of the data set. This can then serve as the basis for an F-test. For either approach, the resulting grid magnitude maps (using 6-fold-symmetry) for all used object groupings were visually indistinguishable from the respective control maps (using 5-fold symmetry) and higher GM values were not found to be confined to any single region of the brain. We therefore abstained from further detailed and ROI-based analysis.

| Set | | Cluster | | | | Peak | | | | | Location | | |
|---|---|---|---|---|---|---|---|---|---|---|---|---|---|
| p | c | $p_{FWE}$ | $p_{FDR}$ | k | $p_{unc}$ | $p_{FWE}$ | $p_{FDR}$ | T | z | $p_{unc}$ | x | y | z |
| **Hierarchy – Iteration** | | | | | | | | | | | | | |
| 0,018 | 2 | 0,003 | 0,005 | 119 | 0,000 | 0,091 | 0,426 | 4,787 | 4,479 | 0,0000 | -45 | 14 | 45 |
| | | | | | | 0,943 | 0,829 | 3,633 | 3,486 | 0,0002 | -45 | 23 | 32 |
| | | | | | | 0,980 | 0,888 | 3,506 | 3,373 | 0,0004 | -33 | 14 | 58 |
| | | 0,000 | 0,001 | 169 | 0,000 | 0,152 | 0,426 | 4,620 | 4,340 | 0,0000 | -63 | -40 | 0 |
| | | | | | | 0,363 | 0,692 | 4,295 | 4,065 | 0,0000 | -60 | -55 | 10 |
| | | | | | | 0,508 | 0,692 | 4,142 | 3,933 | 0,0000 | -54 | -37 | -3 |
| **Interaction 1: Task x Position (2 vs 1) = [H2-H1]-[I2-I1]** | | | | | | | | | | | | | |
| 0,080 | 1 | 0,000 | 0,000 | 209 | 0,000 | 0,000 | 0,000 | 6,241 | 5,622 | 0,0000 | 0 | 35 | 45 |
| **Interaction 2: Task x Position (1 vs 2) = [H1-H2]-[I1-I2]** | | | | | | | | | | | | | |
| 0,000 | 11 | 0,000 | 0,000 | 468 | 0,000 | 0,000 | 0,005 | 6,359 | 5,710 | 0,0000 | 33 | -7 | 61 |
| | | | | | | 0,001 | 0,011 | 5,982 | 5,428 | 0,0000 | 39 | -22 | 39 |
| | | | | | | 0,701 | 0,349 | 3,955 | 3,770 | 0,0001 | 30 | -34 | 48 |
| | | 0,000 | 0,000 | 311 | 0,000 | 0,006 | 0,025 | 5,593 | 5,127 | 0,0000 | -30 | 35 | 32 |
| | | | | | | 0,288 | 0,158 | 4,389 | 4,145 | 0,0000 | -27 | 29 | 39 |
| | | | | | | 0,951 | 0,521 | 3,611 | 3,467 | 0,0003 | -24 | 50 | 10 |
| | | 0,000 | 0,000 | 978 | 0,000 | 0,007 | 0,025 | 5,562 | 5,104 | 0,0000 | 30 | 32 | 26 |
| | | | | | | 0,011 | 0,030 | 5,404 | 4,979 | 0,0000 | 21 | 20 | 10 |
| | | | | | | 0,023 | 0,042 | 5,202 | 4,818 | 0,0000 | -9 | 44 | -9 |
| | | 0,000 | 0,000 | 417 | 0,000 | 0,008 | 0,025 | 5,509 | 5,062 | 0,0000 | -21 | 11 | 4 |
| | | | | | | 0,024 | 0,042 | 5,194 | 4,811 | 0,0000 | -18 | 23 | 0 |
| | | | | | | 0,046 | 0,046 | 4,999 | 4,653 | 0,0000 | -24 | -13 | -19 |
| | | 0,023 | 0,015 | 74 | 0,003 | 0,030 | 0,042 | 5,132 | 4,761 | 0,0000 | 27 | -43 | 64 |
| | | 0,015 | 0,011 | 82 | 0,002 | 0,040 | 0,046 | 5,043 | 4,689 | 0,0000 | 9 | -28 | 36 |
| | | | | | | 0,787 | 0,366 | 3,866 | 3,692 | 0,0001 | 6 | -19 | 42 |
| | | | | | | 0,946 | 0,509 | 3,625 | 3,479 | 0,0003 | -3 | -37 | 45 |
| | | 0,014 | 0,011 | 84 | 0,002 | 0,042 | 0,046 | 5,029 | 4,677 | 0,0000 | 30 | -16 | -19 |
| | | | | | | 0,161 | 0,097 | 4,601 | 4,323 | 0,0000 | 33 | -22 | -12 |
| | | | | | | 0,300 | 0,158 | 4,373 | 4,131 | 0,0000 | 24 | -13 | -12 |
| | | 0,000 | 0,000 | 206 | 0,000 | 0,062 | 0,052 | 4,911 | 4,581 | 0,0000 | -24 | -49 | 64 |
| | | | | | | 0,463 | 0,248 | 4,187 | 3,972 | 0,0000 | -6 | -55 | 55 |
| | | | | | | 0,792 | 0,366 | 3,860 | 3,687 | 0,0001 | 3 | -46 | 55 |
| | | 0,028 | 0,017 | 70 | 0,004 | 0,075 | 0,057 | 4,852 | 4,532 | 0,0000 | -27 | -49 | -28 |
| | | | | | | 0,738 | 0,365 | 3,918 | 3,738 | 0,0001 | -18 | -52 | -25 |
| | | | | | | 0,991 | 0,690 | 3,435 | 3,309 | 0,0005 | -12 | -55 | -19 |
| | | 0,000 | 0,000 | 309 | 0,000 | 0,100 | 0,068 | 4,757 | 4,454 | 0,0000 | -24 | -13 | 58 |
| | | | | | | 0,341 | 0,180 | 4,321 | 4,087 | 0,0000 | -36 | -7 | 58 |
| | | | | | | 0,588 | 0,300 | 4,065 | 3,866 | 0,0001 | -24 | 2 | 58 |
| | | 0,003 | 0,003 | 116 | 0,000 | 0,170 | 0,099 | 4,581 | 4,307 | 0,0000 | 54 | -4 | -25 |
| | | | | | | 0,505 | 0,260 | 4,145 | 3,936 | 0,0000 | 45 | 2 | -35 |
| | | | | | | 0,630 | 0,328 | 4,025 | 3,831 | 0,0001 | 51 | -16 | -16 |

**Table S1. SPM statistics summary table for the TASK x POSITION model.**

| | H | cn | k | %c | nvcr | %r | x | y | z | *Z-score* |
|---|---|---|---|---|---|---|---|---|---|---|
| Interaction Task x Experience (Late vs Early) *[HL-HE]-[IL-IE]* | | | | | | | | | | |
| Frontal Inf Oper | R | 1 | 75 | 67 | 50 | 13 | 57 | 11 | 13 | 5.10 |
| Precentral | L | 2 | 98 | 39 | 38 | 4 | -27 | -1 | 42 | 4.61 |
| Temporal Mid | R | 3 | 65 | 82 | 53 | 4 | 54 | -67 | 13 | 4.51 |
| Frontal Sup 2 | R | 4 | 159 | 52 | 83 | 6 | 33 | -7 | 61 | 4.29 |
| Supp Motor Area | R | 5 | 104 | 59 | 61 | 9 | -9 | 5 | 45 | 3.89 |

**Table S2. Summary of whole brain results for the main effect of TASK and interaction TASK x EXPERIENCE.**

Significant activations were found for the contrast [HL-HE]-[IL-IE] but not for [HE-HL]-[IE-IL]. All activations are significant at cluster-level FWE-corrected $p$ <.05, and voxel-level at $p$<.001. Cluster and main peaks were labeled using the AAL3 toolbox. In this summary, only activations are included that make up at least 5% of a cluster and of the labeled area. A minimum of one area per cluster was displayed irrespective of that. Voxel numbers are based on a voxel size of 3x3x3. Abbreviations: cn – cluster number, k – number of voxels in cluster %c – percentage of the cluster belonging to the area, nv – number of voxels of that cluster within the respective area, %r percentage of region occupied by voxels from the cluster.

| Set | | Cluster | | | | Peak | | | | | Location | | |
|---|---|---|---|---|---|---|---|---|---|---|---|---|---|
| p | c | $p_{FWE}$ | $p_{FDR}$ | k | $p_{unc}$ | $p_{FWE}$ | $p_{FDR}$ | T | z | $p_{unc}$ | x | y | z |
| **Interaction: Task x Experience = [HL-HE]-[IL-IE]** | | | | | | | | | | | | | |
| 0,000 | 5 | 0,024 | 0,040 | 75 | 0,003 | 0,006 | 0,040 | 5,281 | 5,097 | 0,0000 | 57 | 11 | 13 |
| | | | | | | 0,009 | 0,040 | 5,193 | 5,018 | 0,0000 | 51 | 8 | 6,8 |
| | | | | | | 0,954 | 0,773 | 3,502 | 3,444 | 0,0003 | 39 | 2 | 10 |
| | | 0,008 | 0,017 | 98 | 0,001 | 0,050 | 0,096 | 4,745 | 4,609 | 0,0000 | -27 | -1 | 42 |
| | | | | | | 0,493 | 0,409 | 4,012 | 3,927 | 0,0000 | -15 | -7 | 74 |
| | | | | | | 0,787 | 0,495 | 3,745 | 3,675 | 0,0001 | -24 | -13 | 55 |
| | | 0,040 | 0,054 | 65 | 0,005 | 0,076 | 0,121 | 4,634 | 4,506 | 0,0000 | 54 | -67 | 13 |
| | | | | | | 0,116 | 0,164 | 4,511 | 4,393 | 0,0000 | 57 | -61 | 7 |
| | | | | | | 0,624 | 0,409 | 3,897 | 3,818 | 0,0001 | 57 | -55 | -6 |
| | | 0,001 | 0,004 | 159 | 0,000 | 0,168 | 0,213 | 4,400 | 4,290 | 0,0000 | 33 | -7 | 61 |
| | | | | | | 0,371 | 0,320 | 4,130 | 4,038 | 0,0000 | 39 | -1 | 58 |
| | | | | | | 0,642 | 0,409 | 3,881 | 3,803 | 0,0001 | 21 | -7 | 58 |
| | | 0,006 | 0,017 | 104 | 0,001 | 0,538 | 0,409 | 3,972 | 3,889 | 0,0001 | -9 | 5 | 45 |
| | | | | | | 0,571 | 0,409 | 3,943 | 3,862 | 0,0001 | 6 | -4 | 58 |
| | | | | | | 0,754 | 0,464 | 3,779 | 3,707 | 0,0001 | 6 | 5 | 52 |

**Table S2.** SPM statistics summary table for the TASK x EXPERIENCE model.
